# The histone methyltransferase SDG26 shapes cold stress responses in Arabidopsis through chromatin-based regulation of ABA-dependent and ABA-independent pathways

**DOI:** 10.1101/2025.11.25.690385

**Authors:** Xue Zhang, Julie Zumsteg, Mathieu Erhardt, Wen-Hui Shen, Alexandre Berr

## Abstract

Plants constantly face adverse environmental conditions, including temperature drops that can severely impair growth and productivity. To cope with such stresses, they have evolved complex mechanisms of transcriptional reprogramming. While various cold-responsive pathways have been described, the contribution of chromatin-level regulation, and in particular histone modifications, remains largely obscure. Here, we identify the histone methyltransferase SET DOMAIN GROUP 26 (SDG26) as a positive regulator of cold stress responses in *Arabidopsis thaliana*. We show that *SDG26* is transcriptionally induced and post-transcriptionally stabilized by cold, and that its loss of function leads to increased freezing tolerance but reduced drought tolerance. At the molecular level, SDG26 promotes expression of cold-responsive genes, including members of the CBF-COR regulon, through direct binding and histone H3 lysine 36 trimethylation (H3K36me3) at their chromatin. Concomitantly, SDG26 modulates abscisic acid (ABA) biosynthesis, catabolism, and transport, thereby promoting ABA accumulation, stomatal closure, and drought tolerance. Collectively, our results reveal that SDG26 integrates ABA-dependent and ABA-independent pathways to fine-tune Arabidopsis responses to abiotic stresses. We further establish SDG26 as a chromatin modifier contributing to stress-responsive H3K36me3 enrichment at specific loci. Together, our work identifies SDG26 as a chromatin-based hub balancing cold acclimation with water conservation, thereby enhancing plant resilience.

## Introduction

Plants, as sessile organisms, must continuously adapt to fluctuating environmental conditions and withstand diverse stressors throughout their life cycle. Exposure to these stressors elicits extensive physiological, biochemical, and molecular changes that can severely constrain plant growth and pose a threat to agricultural productivity (Zhang *et al*., 2022). Epigenetic mechanisms, which provide more rapid and reversible responses compared to genetic mutations, have been notably implicated in environmental stress responses, adaptation, and memory in plants (Gaude *et al*., 2024). Among them, histone lysine methylation has emerged as a central regulatory axis that modulates chromatin accessibility. Histone lysine methylation is catalyzed by histone lysine methyltransferases (HKMTs), which do not directly alter chromatin conformations but instead serves as a recruitment platform for specific “readers” that shape transcriptional outcomes in a context-dependent manner (Hu and Du, 2022). Depending on the lysine residue targeted and the number of methyl groups added (mono-, di-, or tri-methyl), this versatile post-translational modification can exert dual effects on gene transcription, either activating or repressing it. In *Arabidopsis thaliana*, about 40 putative HKMTs have been identified and grouped into at least five classes based on sequence homology and domain organization (Springer *et al*., 2003). The SET DOMAIN GROUP (SDG) family includes ABSENT, SMALL OR HOMEOTIC DISCS1 (ASH1) and TRITHORAX (TRX)-related enzymes that typically associate with transcriptional activation via H3K4 and/or H3K36 methylation. Within this family, SDG26/ASHH1 has been studied primarily for its role in flowering-time control. Unlike mutants for its homologs SDG8/ASHH2 (Zhao *et al*., 2005) or *SDG7/ASHH3* (Lee *et al*., 2015), as well as other TRX subfamily members such as *ATX1/SDG27* (Pien *et al*., 2008), *SDG2/ATRX3* (Guo *et al*., 2010; Berr et al., 2010), and *SDG25/ATXR7* (Berr *et al*., 2009)(Tamada *et al*., 2009), the *sdg26* mutant exhibits a unique late-flowering phenotype. Notably, even *CURLY LEAF* (*CLF*), a member of the repressive E(z) family, produces an early-flowering phenotype when mutated (Jiang *et al*., 2008).

SDG8 delays flowering by activating the transcription of the floral repressor *FLOWERING LOCUS C* (*FLC*) through H3K36 di- and tri-methylation (Xu *et al*., 2008). In contrast, SDG26 participates in a multi-subunit complex involving autonomous pathway components such as the H3K4 demethylase FLOWERING LOCUS D (FLD), the transcriptional repressor LUMINIDEPENDENS (LD), ELF4-like proteins (EFL2 and EFL4), and the yeast COMPASS complex subunit SWD2 homolog APRF1/S2La (Fang *et al*., 2020)(Qi *et al*., 2022). Within this complex, SDG26 links RNA 3′-end processing to chromatin regulation, repressing *FLC* transcription (Xu *et al*., 2021). Beyond this repressive function, SDG26 also contributes to transcriptional activation by promoting the expression of the floral integrator *SUPPRESSOR OF OVEREXPRESSION OF CONSTANS1* (*SOC1*) through H3K4 and H3K36 trimethylation (Berr *et al*., 2015)(Bing Liu *et al*., 2016a). These dual activities suggest that SDG26 is a versatile chromatin modifier that can act either as a repressor or activator depending on genomic context and/or interacting partners.

Although most studies of SDG26 have focused on floral transition, emerging evidence indicates that its functions extend further. SDG26 has been implicated in the regulation of pre-messenger RNA splicing (Pajoro *et al*., 2017), trichome growth and development (Zeng *et al*., 2023), and the DNA damage response through mechanisms that remain to be defined (Campi *et al*., 2012)(Roitinger *et al*., 2015). Notably, previous transcriptomic analysis of *sdg26* mutants revealed that nearly one-third of differentially expressed genes are linked to stress responses (Boyu Liu *et al*., 2016)(Berr *et al*., 2015). *In silico* analysis suggests that SDG26 expression is stress-responsive to cold, drought, and salt (Naika *et al*., 2013). Moreover, its promoter contains cis-regulatory elements such as putative W-box motifs, which are binding sites for WRKY transcription factors that mediate numerous biotic and abiotic stress responses (Phukan *et al*., 2016). Together, these observations raised the possibility that SDG26 functions beyond development, contributing to stress adaptation.

In this study, we demonstrate that SDG26 is a critical regulator of Arabidopsis responses to cold stress. We show that *SDG26* transcription is induced by cold, salt, and drought stress, and that its protein abundance is stabilized under cold conditions. At the molecular level, we reveal that *SDG26* modulates a subset of cold-responsive genes through both CBF-COR pathway and ABA homeostasis. Loss of *SDG26* function impairs ABA accumulation, delays cold-induced stomatal closure, and reduces seed dormancy, while paradoxically enhancing freezing tolerance but compromising drought resistance. Collectively, our findings establish SDG26 as a chromatin-based integrator of ABA-dependent and ABA-independent pathways, highlighting its role in fine-tuning stress adaptation and in coordinating responses to multiple abiotic challenges.

## Materials and Methods

### Plant materials

All lines used in this study are derived from the Columbia ecotype (Col0) background of *Arabidopsis thaliana*. Col0 is referred to as the wild-type (WT) plant. The mutants *sdg26-1* (Xu *et al*., 2008) and *soc1-2* (Lee *et al*., 2000) have been previously described. The *sdg26 soc1* double mutant and all the transgenic lines (*pSDG26-GUS*, *p+gSDG26-myc* and *p35S+gSDG26-GFP*) mentioned herein after were generated through genetic crosses or Agrobacterium-mediated transformation. Homozygotes were characterized by antibiotic resistance selection and/or PCR-based genotyping using specific primers (Supplementary Table S2).

### Generation of transgenic lines

The 1772 bp *SDG26* promoter region, the *SDG26* gene along with its promoter region, and the SDG26 CDS were separately PCR amplified using specific primer pairs (Supplementary Table S2). These were then inserted into the Gateway donor vector pDONR207 and subcloned into the Gateway binary vectors pGWB633, pGWB19 and pB7FWG2.0. The resulting constructs (*i.e*., *pSDG26-GUS*, *p+gSDG26-MYC* and *p35S+gSDG26-GFP*) were introduced into *Agrobacterium tumefaciens* GV3101, and transformation was carried out by floral dip. T2 homozygous transgenic Arabidopsis plants were obtained through antibiotic resistance selection.

### *In situ* hybridization and GUS staining

Digoxigenin labeling of RNA probes, tissue preparation, and in situ hybridization were performed as previously described (Berr *et al*., 2010). Tissue sections were 8 μm thick. A 214 bp fragment covering the 3’-end and the 3’-UTR of *SGD26* was obtained through PCR amplification using specific primers (see Table S1). This fragment was cloned into the pGEM-T Easy vector (Promega) and used for the preparation of sense and antisense probes. For GUS expression analysis via histochemical staining, the T2 progeny from six independent homozygous primary transformants were examined as previously described (Berr, *et al*., 2010).

### Plant micrografting

Micrografting between the hypocotyls of rootstocks and scions was carried out without collars on 6-day-old seedlings, as previously described (Regnault *et al*., 2015). Successful grafts were then transferred to MS medium for fluorescence microscopic observation or to soil for flowering time analyses.

### Stress treatments

Plants grown for 10 days on Murashige and Skoog (MS) medium in a growth chamber under long-day (LD; 16 h light: 8 h dark) photoperiod at 21°C were subjected to various types of stress. For chemical treatments, plants grown *in vitro* were transferred to liquid MS medium supplemented with either 100 mM NaCl, 10 μM ABA (Sigma-Aldrich, A1049), 10 μM ACC (Sigma-Aldrich, A3903), 1 μM Kinetin (Sigma-Aldrich, K3378), 1mM SA (Sigma-Aldrich, S5922), or 10 μM MeJA (Sigma-Aldrich, 392707). For cold stress, plates containing plants were transferred to a 4°C cold chamber for 4 or 8h. For drought stress, plants grown in soil under medium-day (MD; 12 h light: 12 h dark) photoperiod at 21°C for 10 days were either watered or left unwatered. After a week, the stressed plants were re-watered for 3 days, and their survival rate were recorded. In parallel, water loss was measured using detached leaves from 10-day-old non-stressed plants by weighting at different time points. Finally, freezing tolerance was assessed in plants grown for 2 weeks in soil at 21°C by exposing them to -10°C. After 2 hours, the plants were allowed to recover in a growth chamber at 21°C, and survival rates were evaluated after one week.

### Germination tests

Seeds were surface-sterilized 5 min in 75% ethanol with 0.01% (v/v) Triton-X 100, rinsed with sterile distilled water and distributed on the surface of ½ MS plates in Petri dishes. Germination was assessed by counting the visible radicle emergence of seeds subjected or not to cold pretreatment (i.e., stratification for 48h at 4°C in the dark). Five biological replicates consisting of 100 seeds per replicate were used for each experiment.

### Hormones quantification

ABA and jasmonate were identified and quantified using UPLC–MS/MS, as described previously (Smirnova *et al*., 2017). Briefly, Col0, *sdg26-1*, *soc1-2*, and *sdg26soc1* plants were grown on MS for two weeks under LD conditions and subjected to cold stress at 4°C or maintained at 21°C. Approximately 150 mg of fresh plant material from each genotype was collected, ground in an extraction solution (6xV/W; 70% methanol, 29% water, 1% acetic acid) containing internal standard (deuterated ABA or dABA), and immediately frozen in liquid nitrogen. The extraction solutions were analyzed using the following MRM transitions: JA (209 > 59, -), JA-Ile (324 > 151, +), ABA (263 > 153, -), and dABA (269 > 159, -), with + and - indicating analysis in positive or negative mode, respectively. Quantitative profiles were obtained using an EVOQ Elite LC-TQ (Bruker) equipped with an electrospray ionization source and coupled to a Dionex UltiMate 3000 UHPLC system (Thermo). Data acquisition was performed with the MS Workstation 8 for mass spectrometry, while liquid chromatography was controlled by Bruker Compass Hystar 4.1 SR1 software. Data analysis was conducted using the MS Data Review software. Absolute quantifications were achieved by comparing sample signals using dose-response curves established with pure compounds.

### Stomata analysis

Epidermal strips from 2-week-old plants were fixed in a solution containing 1% glutaraldehyde, 10 mM PIPES (pH 7.0), 5 mM MgCl_2_ and 5 mM EGTA for 1 hour. The samples were then cleared overnight in chloral hydrate, and stomata were examined under a Nikon Eclipse 800 microscope. Microscopy images were captured using a Hitachi S-3400N (Hitachi High-Technologies Europe). All images were processed using ImageJ (NIH).

### Western blotting

Western blot analysis was performed on total protein extracts prepared from 2-week-old seedlings, as described previously (Zhang *et al*., 2020). Protein samples were separated on 8% SDS-PAGE and transferred onto Immobilon-P PVDF membranes (Merck Millipore). The membranes were probed with antibodies against GFP (11120; Sigma-Aldrich), MYC (C3956; Sigma-Aldrich), or H3 (ab12079; Abcam), followed by a goat anti-mouse or anti-rabbit HRP-conjugated secondary antibodies (62-6520, Thermofisher; A9169, Sigma-Aldrich). Chemiluminescence was detected with ECL+ (Lumi-Light PLUS Western Blotting; Roche) and visualized with ECL films (Bio-Rad). The intensity of individual bands was quantified using ImageJ (NIH).

### Gene expression analyses by qPCR

Total RNA was isolated using the Nucleospin RNA Plant Kit (Macherey-Nagel). First-strand cDNA was synthesized using SuperScript® IV Reverse Transcriptase (Invitrogen) with an oligo-dT primer, following the manufacturer’s recommendations. Transcript abundance was quantified in triplicates using gene-specific primers (Supplementary Table S2) in a 10 µL reaction mix containing Takyon™ No Rox SYBR® MasterMix Blue dTTP on a LightCycler 480 (Roche). *EXP* (AT4G26410) and *TIP4.1* (AT4G34270) were used for normalization. Relative expression levels were calculated using the 2^−ΔΔCT^ method.

### Histone modifications analyses

Chromatin immunoprecipitation was performed as previously described (Boyu Liu *et al*., 2016). Chromatin was precipitated using anti-H3 (ab12079; Abcam), anti-trimethyl-H3K36 (ab9050; Abcam), anti-MYC (C3956; Sigma-Aldrich), anti-GFP (ab290; Abcam) or anti-SDG26 (Berr *et al*., 2015), along with protein A magnetic beads (Magna-ChIP, Millipore). After immunoprecipitation, DNA was recovered using the NucleoSpin Gel and PCR Clean-up kit (Macherey-Nagel) and analyzed by real-time PCR (LightCycler 480II; Roche) in conjunction with the SYBR Green Master mix using gene-specific primers (Supplementary Table S2).

### Statistical methods

Statistical analyses were performed in R (http://www.r-project.org/) using a Student’s *t*-test with Benjamini–Hochberg false discovery rate (FDR) correction.

## Results

### *SDG26* is ubiquitously expressed in Arabidopsis

To gain insight into *SDG26* functions, we first examined its expression pattern. The activity of the *SDG26* promoter was analyzed using a β-glucuronidase (GUS) reporter lines (pSDG26-GUS). Histochemical staining of independent homozygous transformants consistently revealed broad expression across multiple organs and developmental stages. In adult plants, strong GUS activity was detected in leaf tissues (Figure 1A) and at organ-stem junctions (Figure 1B-D). In root, expression was absent from the root cap but prominent in the vascular system, pericycle, and endodermis at the meristematic zone, as well as at the base of emerging lateral root primordia (Figure 1E). In seedlings, GUS activity was observed in the vasculature of roots, leaves, and cotyledons, as well as at the leaf axillary meristem (Figure 1F-H). Consistent with these findings, *in situ* hybridization showed strong transcript accumulation in young and actively dividing tissues, including the shoot and floral apical meristems, developing embryos, pollen, and ovule primordia. The labeling was heterogeneous, with clusters of highly stained cells (Figure 2A-G). RT-qPCR analysis further confirmed ubiquitous expression of *SDG26* in wild-type plants, with relatively stable transcript levels throughout the diurnal cycle (Figure 2H and I). To investigate subcellular localization, we generated *sdg26-1* lines expressing a C-terminal GFP-tagged SDG26 under the 35S promoter (*sdg26-1* p35S::SDG26-GFP). These plants complemented the late-flowering phenotype of *sdg26-1*, indicating functional rescue. Confocal microscopy revealed nuclear localization of SDG26-GFP, with exclusion from the nucleolus (Supplementary Figure 1A). In addition, grafting experiments demonstrated that SDG26 does not exhibit long-distance mobility between scion and rootstock (Supplementary Figure 1B). Together, these data indicate that SDG26 is a widely expressed nuclear protein, consistent with the broad expression domains reported for other Trithorax group (TrxG) methyltransferases such as *SDG8* (Berr, *et al*., 2010) and *ATX1* (Saleh *et al*., 2007). This overlap suggests potential functional redundancy and/or mutual interactions among TrxG proteins (Valencia-Morales *et al*., 2012).

**Figure 1:**
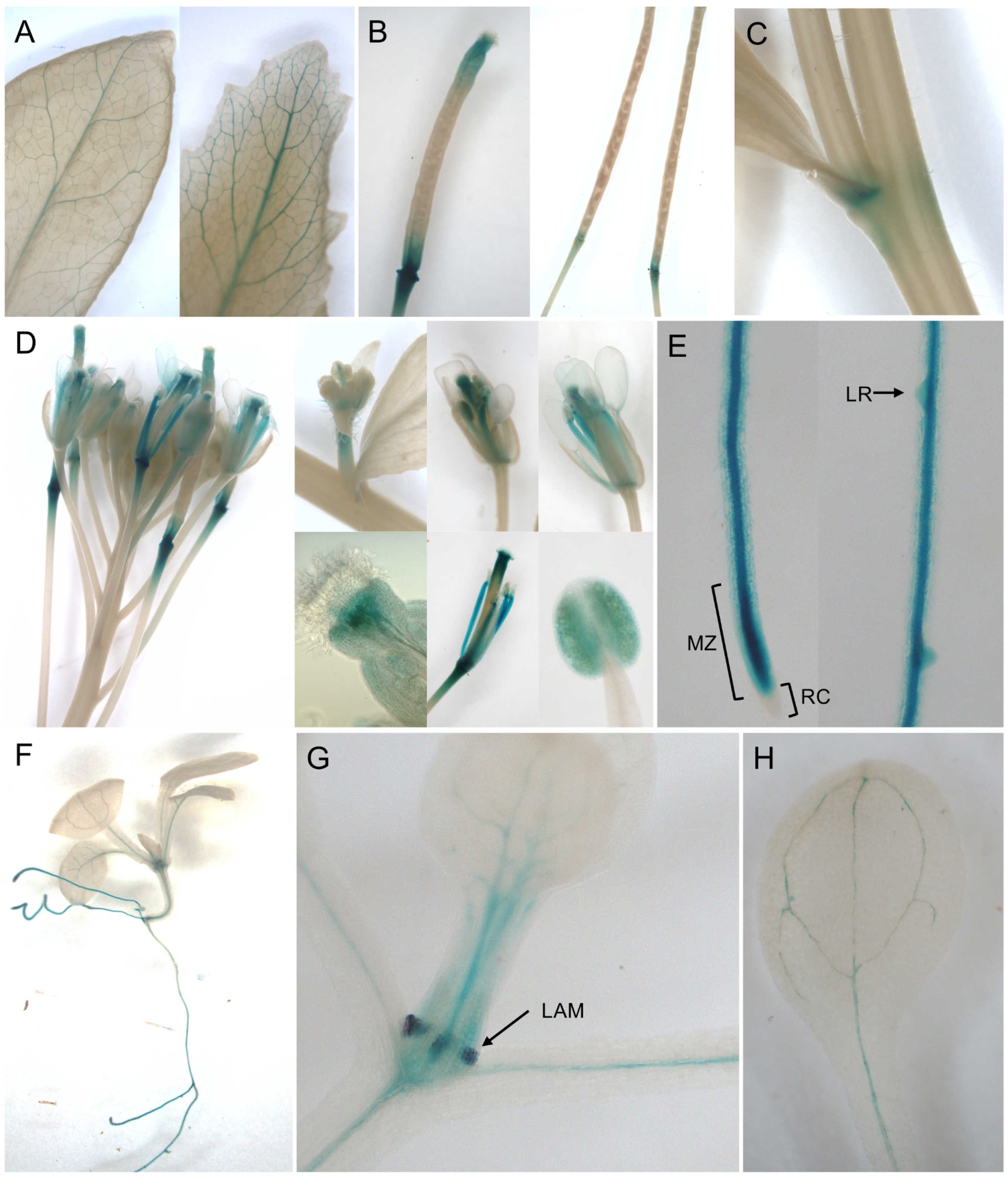
GUS staining in transgenic plants harboring a GUS reporter gene driven by the SDG26 promoter. (**A**) Mature (left) and cauline (right) leaves from 6-week-old plants. (**B**) Siliques at different developmental stages, 3 to 4 Days Post Anthesis (DPA; left) and 9 to 11 DPA. (**C**) Base of shoot branches and cauline leaf petiole. (**D**) Total inflorescence and different flower details, from young flower buds (top left), young flower (top middle), open flower (top right), stigma after pollination (bottom left), young silique (bottom middle) and anther (bottom right). (**E**) Root tip with the root cap (RC) and the meristematic zone (MZ; left) and maturation zone with emerging lateral root (LR; right). (**F**) 15-day-old seedling. (**G**) Closeup of the lateral leaf axillary meristem (LAM) of a 10-day-old seedling. (**H**) Cotyledon of a 10-day-old seedling.

**Figure 2:**
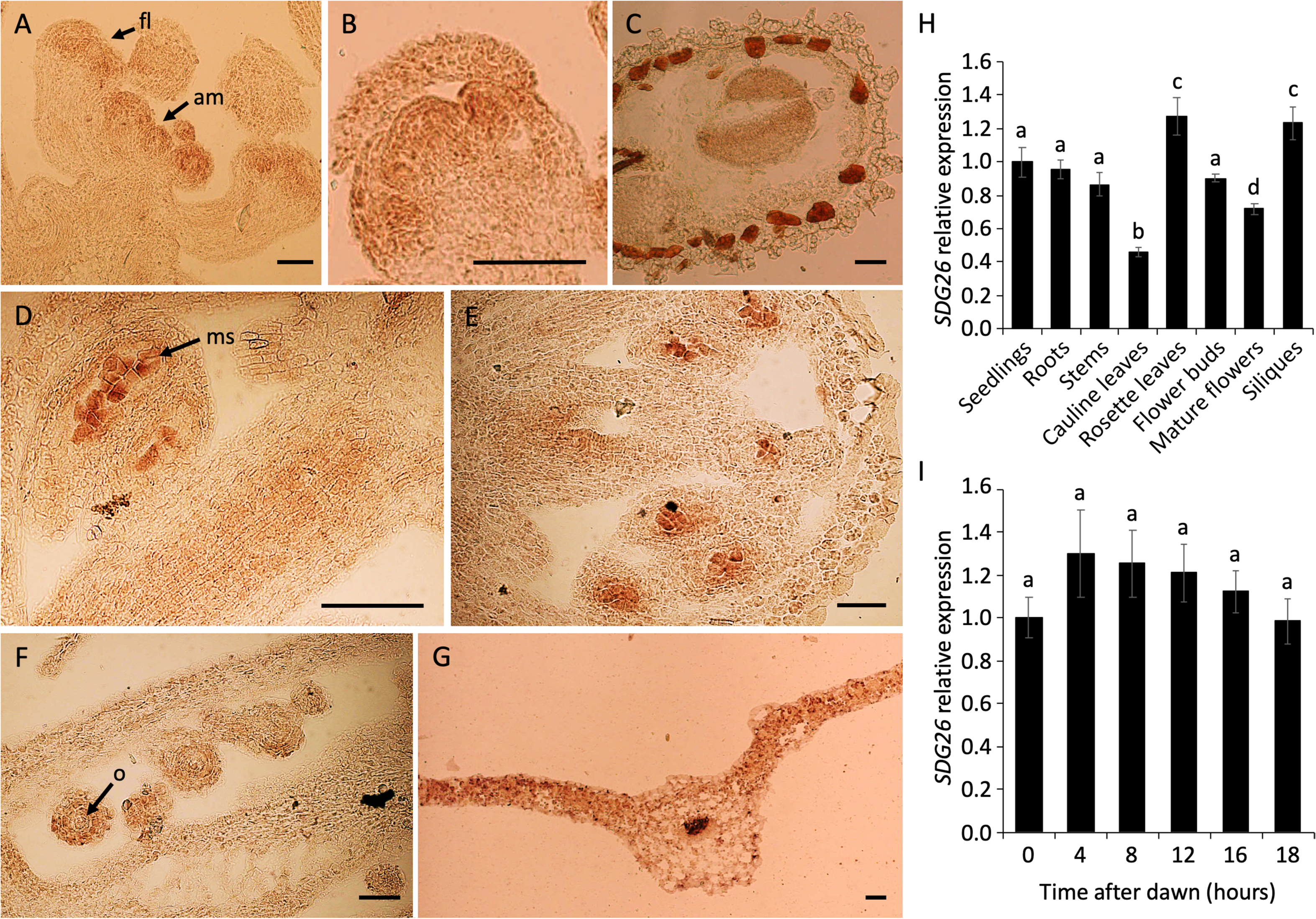
Expression pattern analysis of *SDG26* by *in situ* hybridization and qPCR. A-G) *In situ* hybridization analysis of *SDG26* in: (**C**) a floral apex with (fl) for flower primordia and (am) for apical meristem; (**D**) a vegetative apex with the Shoot Apical Meristem (SAM) in the center; (**E**) an embryo at mid-torpedo stage; (**F** and **G**) anthers at stage 9 with (ms) for microspore; (**H**) a gynoecium with (o) for ovule; and (I) a leaf dissected from a 15-day-old seedling with at the center the mid-vein. Bars = 100 µm. Real-time quantitative PCR (qPCR) analysis of *SDG26* (**H**) in different tissues of Arabidopsis wild type plants and (**I**) in 10-day-old seedlings grown under mid-day (MD) conditions (16h light/8h dark). Transcript levels are expressed relative to the transcript level in seedlings for **H** or at time 0h after dawn for **I**. Values are the means ±SD of at least three biological replicates. Letters indicate significant differences (Student’s t-test with Benjamini-Hochberg FDR correction, *P* < 0.05).

### *SDG26* is induced by different abiotic stresses but not by stress-related phytohormones

Because the SDG26 promoter contains a putative W-box motif and previous transcriptomic data suggested stress responsiveness, we next tested whether *SDG26* is induced by abiotic stress. RT-qPCR analysis showed that cold, drought, and salt treatments each triggered a modest but significant transient increase in *SDG26* transcript abundance (Figure 3A-C). By contrast, no significant induction was observed upon treatment with stress-related signaling molecules such as ABA, ACC, kinetin, salicylic acid, or methyl jasmonate (Supplementary Figure 3A-E). To validate these results, we monitored GUS activity in the pSDG26-GUS line above described. Cold treatment enhanced GUS staining compared with control conditions (Figure 3D), confirming our TR-qPCR results. We then tested whether transcript accumulation translated into protein-level changes. To this end, we generated *sdg26-1* transgenic plants expressing SDG26 fused to a 10x c-Myc tag under the control of its native promoter (p+gSDG26-MYC; Supplementary Figure 1). In these plants, immunoblot analysis revealed a gradual increase of SDG26 protein during cold exposure (Figure 3E). Importantly, subcellular localization remained unchanged, SDG26-GFP localized exclusively to the nucleus under both control and cold stress conditions (Supplementary Figure 4). Together, these results show that *SDG26* is induced by multiple abiotic stresses at both the transcription and protein levels, but not directly by stress-related phytohormones. This suggests that SDG26 functions as an upstream chromatin regulator responding primarily to environmental cues rather than to hormonal signals.

**Figure 3:**
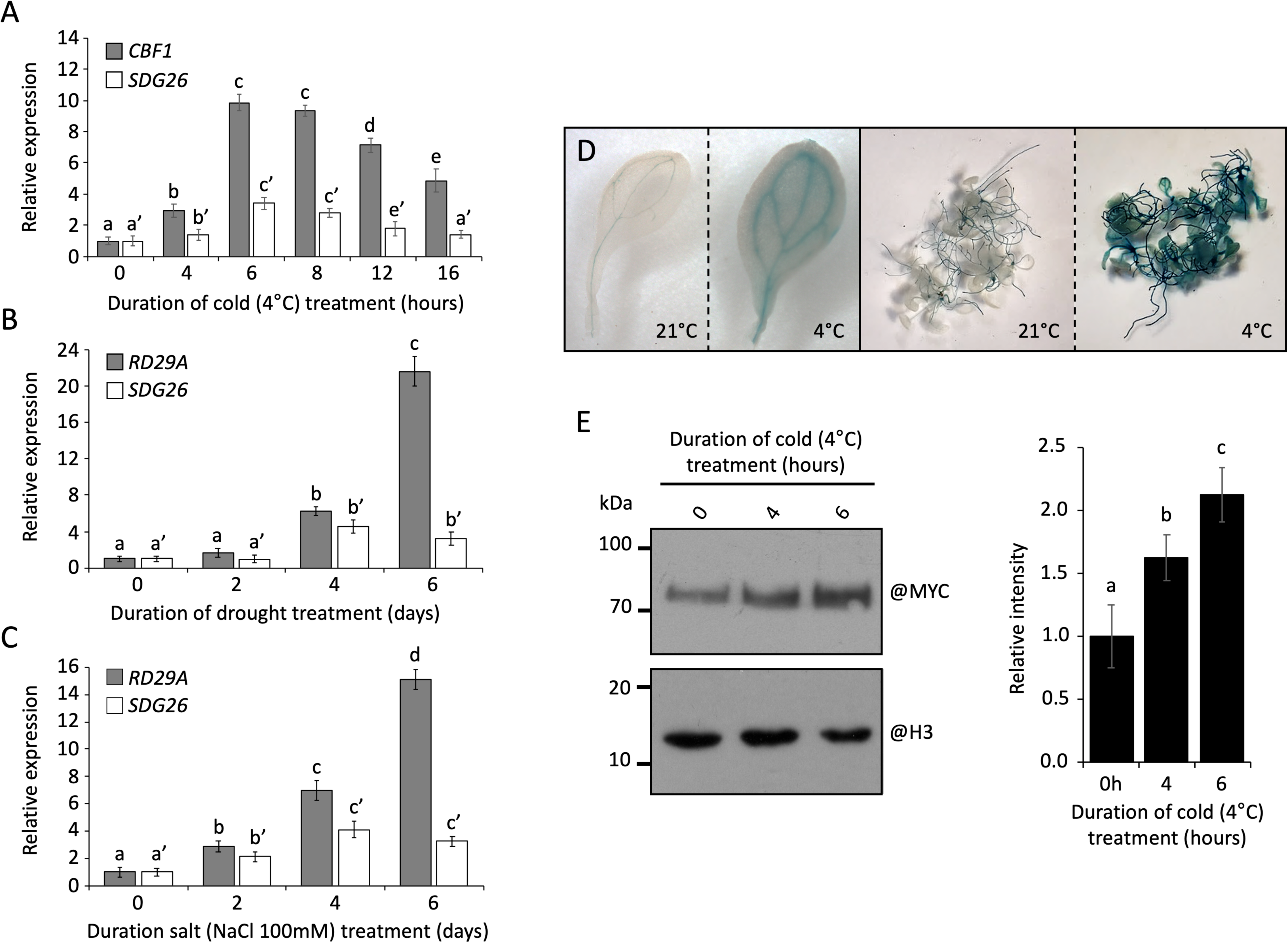
*SDG26* expression analysis in response to different abiotic stresses. The *SDG26* transcript level was quantified in wild-type Col0 plantlets transferred to 4°C (**A**), not watered (**B**) or watered with a salt solution (**C**; NaCl 100mM). For each condition, a gene know to be transcriptionally induced was used as a control. Expression levels of each gene are presented relative to their level before treatment (set as 1) as mean ±SD of at least three biological replicates. Letters indicate significant differences (*P* < 0.05). (**D**) Plantlets expressing a *GUS* reporter gene driven by the *SDG26* promoter (pSDG26-GUS) were transferred to 4°C for 4h. Stressed (4°C) and non-stressed (22°C) transgenic lines were similarly fixed in 80% acetone and stained at 37°C for 6h. (**E)** The level of the SDG26-Myc transgenic protein was measured by Western blot in the *sdg26* transgenic line expressing *p+gSDG26-Myc* before and during cold stress using an anti-Myc antibody. Histone H3 was used as a loading control. Densitometry values ±SD were normalized to H3 and shown relative to time 0 (set as 1). Letters indicate significant differences (*P* < 0.05).

### SDG26 directly binds cold-related genes and regulates their transcription

SDG26 was previously shown to directly regulate *SOC1* expression in flowering-time control (Berr *et al*., 2015). Because *SOC1* also participates in a feedback loop linking flowering and cold response (Seo *et al*., 2009), we examined its regulation by SDG26 during cold stress. Under control conditions (21°C), *SOC1* expression remained relatively stable, while upon cold exposure (4°C), *SOC1* transcript levels gradually increased in both Col0 and *sdg26-1* plants, as well as in our p+gSDG26-Myc transgenic line (Figure 4A; Supplementary Figure 5). Interestingly, these increases were diminished in *sdg26-1* compared to Col0. To investigate how *SDG26* regulates *SOC1* expression, we performed ChIP-qPCR to assess changes in H3K4me3 and H3K36me3 levels, two active histone marks, at the *SOC1* chromatin (Figure 4B-F). As previously reported (Berr *et al*., 2015)(Bing Liu *et al*., 2016b)(Boyu Liu *et al*., 2016), H3K36me3 and H3K4me3 levels were lower in *sdg26-1* compared to Col0 plants grown at 21°C. Upon cold exposure, H3K36me3 levels remained unchanged in both Col0 and *sdg26-1*, whereas H3K4me3 levels increased (Figure 4C and D). To determine whether SDG26 is recruited to SOC1 chromatin in response to cold, we performed ChIP assays using two complementary approaches: (*i*) a monoclonal SDG26 antibody on Col0 and *sdg26-1* chromatin extracts, and (*ii*) a MYC antibody on chromatin extracts from both non-transformed Col0 plants and p+gSDG26-Myc transgenic line. In both cases, SDG26 occupancy at SOC1 chromatin remained unchanged following cold stress (Figure 4E and F). Thus, SDG26 contributes both to *SOC1* basal expression and to its cold-induced upregulation, without changes in recruitment.

**Figure 4:**
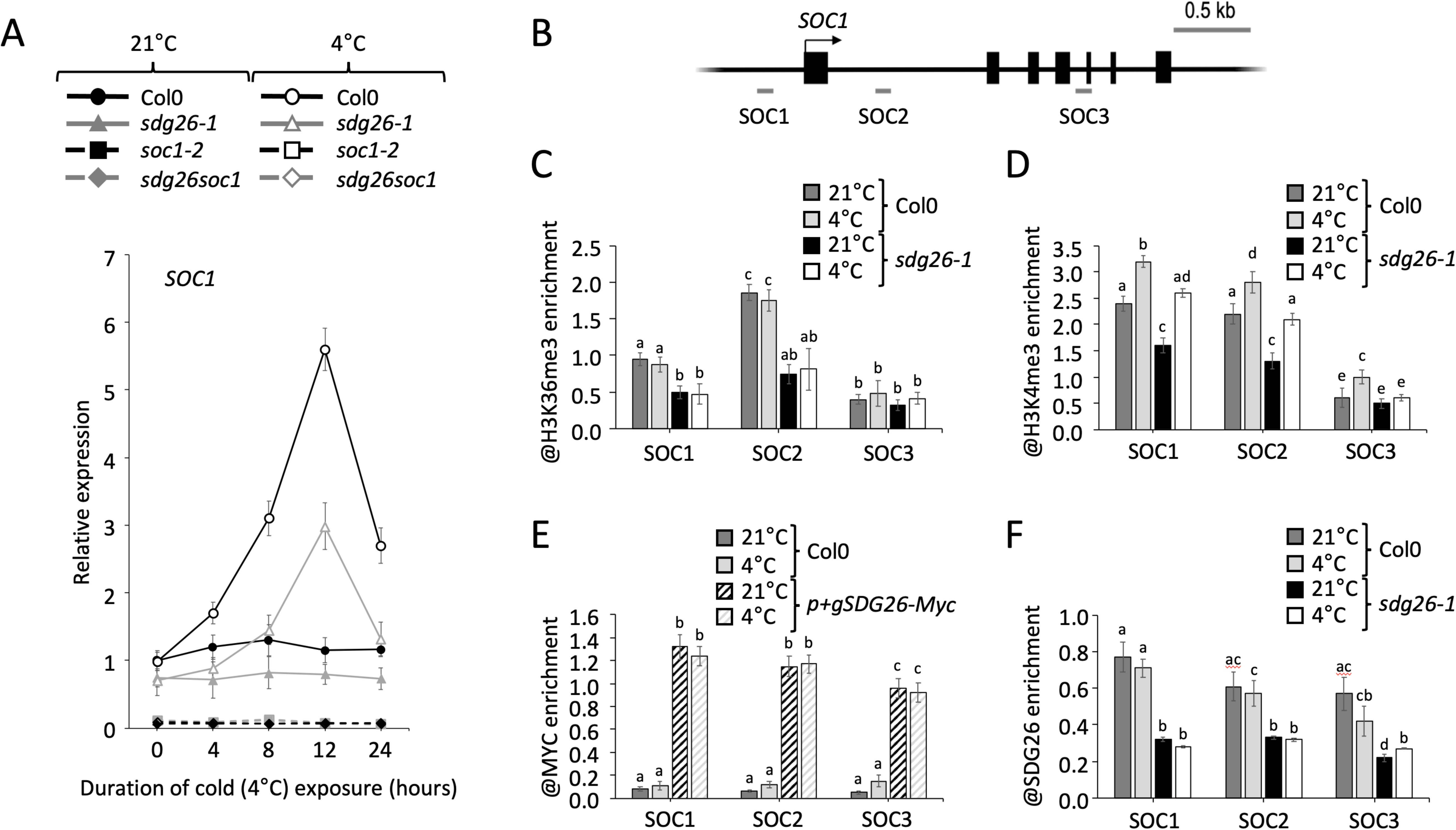
Transcript and chromatin analysis of *SOC1* during cold stress. (**A**) The dynamic expression of *SOC1* was determined in plants treated with a cold stress (4°C) or not (22°C). Data are presented relative to Col0 at 0h (set as 1) as mean ±SD of at least three biological replicates. (**B**) Schematic structure of *SOC1* represented as black boxes for exons and black lines for promoter and introns, with an arrow for the translational start site and amplified regions as grey lines. (**C** and **D**) Levels of H3K36me3 and H3K4me3 along different regions of *SOC1* were measured by ChIP using the same plant material as in (**A**) at 8h. (**E** and **F**) ChIP analysis of SDG26 enrichment at the *SOC1* loci using an anti-SDG26 monoclonal antibody in Col0 or an anti-MYC antibody in *sdg26-1* transgenic line expressing *p+gSDG26-Myc*. Data are presented as the ratio of H3K36me3, H3K4me3, SDG26 and MYC over H3 for each amplified region. Mean values ±SD in (**C** to **F**) are presented based on results from two biological replicates. Letters indicate significant differences (P < 0.05).

SOC1 is a known negative regulator of the cold-responsive CBF genes *CBF1*, *CBF2* and *CBF3*, and their downstream targets, the CBF-target genes *COR15A*, *RD29A*, *KIN1* and *KIN2* (Seo *et al*., 2009). Given that *SDG26* regulates *SOC1* transcription and is itself induced by cold, we next examined whether SDG26 also affects the expression of cold-responsive genes. Consistent with previous report (Seo *et al*., 2009), *CBF* genes and their downstream targets were upregulated in *soc1-2* compared to Col0 under both control and cold conditions (Figure 5A; Supplementary Figure 6A). Surprisingly, despite the lower *SOC1* expression observed in *sdg26-1* and *sdg26soc1* (Figure 4A), *CBF* and its target genes showed weaker induction by cold compared to Col0 (Figure 5A). To resolve this paradox, we tested whether SDG26 directly regulates CBF genes. ChIP-qPCR revealed SDG26 enrichment at CBF1-3 chromatin, which further increased under cold treatment (Figure 5B-D). In parallel, H3K36me3 levels at *CBF1* were reduced in *sdg26-1* relative to Col0 under control conditions and showed impaired increase upon cold exposure (Figure 5E). These results suggest that SDG26 exerts a dual regulatory role. It positively regulates *SOC1*, a repressor of CBFs, while simultaneously directly activating CBF loci through H3K36me3 deposition. This dual action suggests that SDG26 fine-tunes the cold response by balancing repression via SOC1 with direct activation of cold-inducible genes.

**Figure 5:**
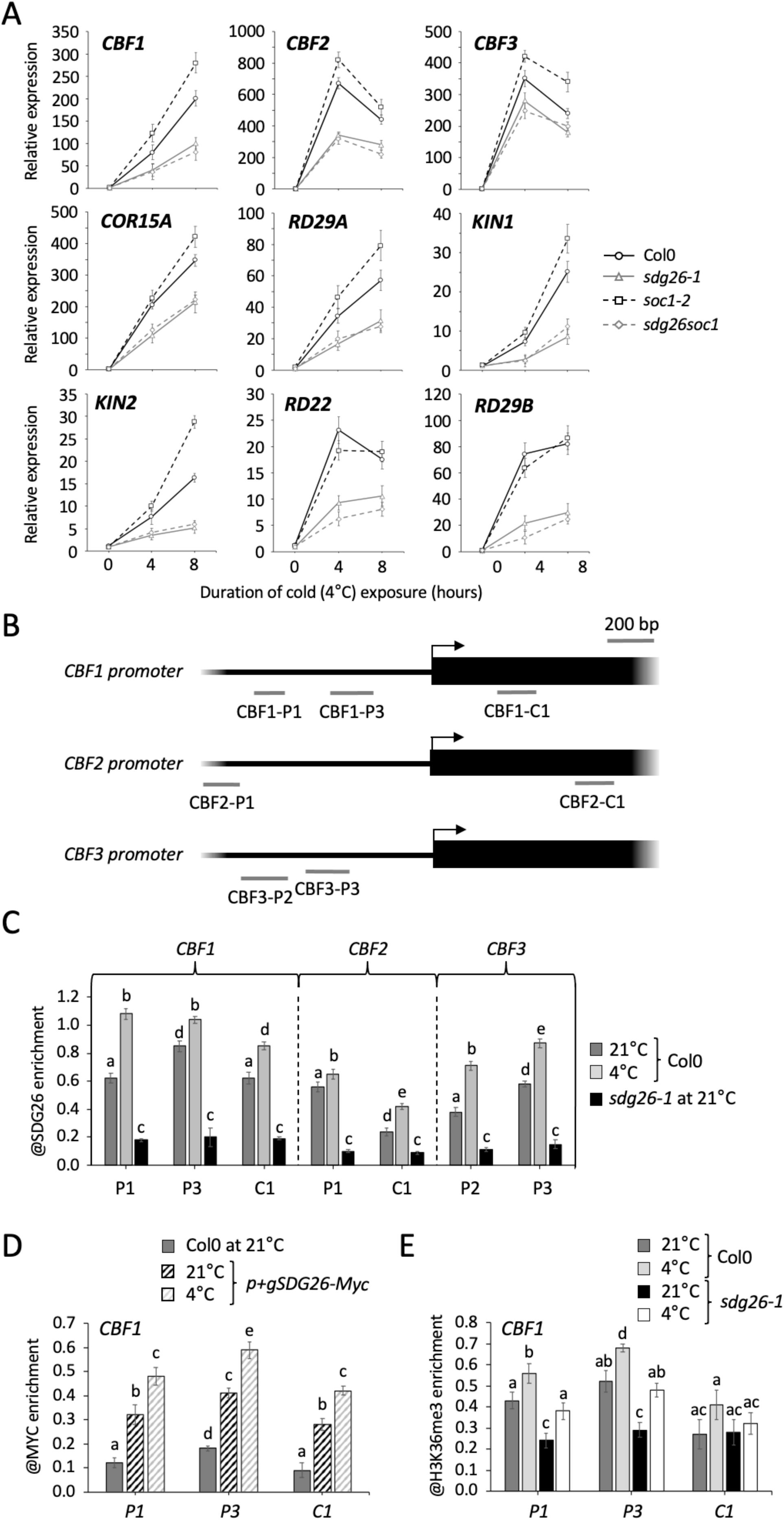
Transcript and chromatin analysis of cold*-*inducible genes during cold stress. (**A**) Dynamic expressions of cold-inducible genes were determined in 2-weeks-old Col0, *sdg26-1*, *soc1-2* and *sdg26soc1* plantlets before (0h) and during (4h and 8h) a cold treatment at 4°C. Data are presented relative to Col0 at 0h (set as 1) as mean ±SD of at least three biological replicates. (**B**) Schematic diagrams adapted from (Shi *et al*., 2012) showing the promoter structures of the *CBF1*, *CBF2* and *CBF3*. The 1.0 kb upstream regions and coding regions are shown, with the translational start site as an arrow and amplified regions as grey lines. (**C** and **D**). Analyses of the SDG26 enrichment at different regions of the *CBF* genes by ChIP using an anti-SDG26 monoclonal antibody in Col0 or an anti-MYC antibody in *sdg26-1 p+gSDG26-Myc*. (**E**) The level of H3K36me3 along the *CBF1* chromatin was measured by ChIP in plants treated with a cold stress (4°C) or not (22°C). For (C, D and E) Data are presented as the ratio of SDG26, SDG26-MYC and H3K36me3 over H3 for each amplified region. Mean values ±SD are presented based on results from two biological replicates.

### SDG26 plays roles in ABA homeostasis

In addition to CBF-dependent pathways, plants activate ABA-dependent but CBF-independent signaling during cold stress (Barnaby *et al*., 2020). Genes such as *RD29B* and *RD22* fall into this category (Zhang *et al*., 2004). Their expression was similarly induced in *soc1-2* and Col0 plants but was markedly reduced in *sdg26-1* and *sdg26soc1* mutants under cold conditions (Figure 5A; Supplementary Figure 6A and B). Given that ABA accumulates in response to cold stress (Pareek *et al*., 2017), this suggested a possible defect in ABA homeostasis. We therefore quantified ABA levels during cold exposure. The phytohormone jasmonate (JA), known to positively regulate cold stress responses (Hu *et al*., 2017), was also quantified. While JA levels remained unchanged across all tested genotypes, ABA accumulation was significantly reduced in *sdg26-1* and *sdg26soc1* compared with Col0 and *soc1-2* at both control and cold conditions (Figure 6A). This indicates that SDG26 regulates cold stress responses, at least in part, by promoting ABA accumulation, which could explain the lower expression of *RD29B* and *RD22* in *sdg26-1* and *sdg26soc1* mutants upon cold exposure.

**Figure 6:**
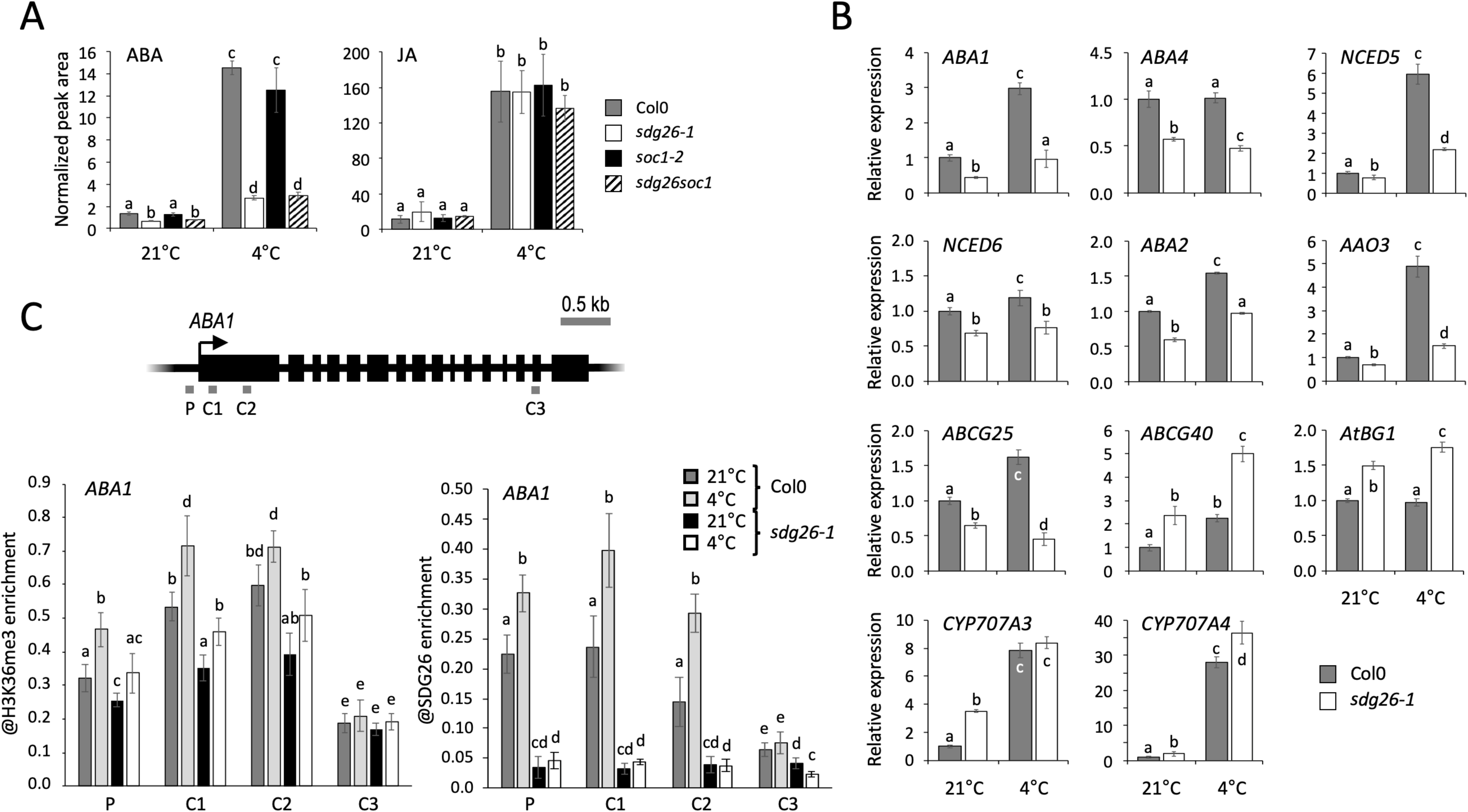
ABA homeostasis in response to cold. (**A**) Quantification of ABA (left) and JA (right) contents were determined in 2-weeks-old Col0, *sdg26-1*, *soc1-2* and *sdg26soc1* plantlets treated with a cold stress (4°C) or not (22°C). Hormonal content is expressed as area of the peaks obtained by UHPLC-MS/MS analysis. (**B**) Expression analyses of key genes involved in regulation of ABA homeostasis in 2-weeks-old Col0 and *sdg26-1* plantlets treated (4°C) or not (RT) with a cold stress at 4°C. Data are expressed relative to Col0 level at RT (set as 1) as mean ±SD of at least three biological replicates. (**C**) The levels of H3K36me3 and SDG26 along different regions of *ABA1* were measured by ChIP in 2-weeks-old plantlets treated (4°C) or not (RT) with a cold stress at 4°C. Data are presented as the ratio of H3K36me3 and SDG26 / H3 for each primer. Mean values ±SD are presented based on results from two biological replicates.

The regulation of ABA biosynthesis, metabolism, and transport is critical for adjusting ABA levels in response to abiotic stress (Baron *et al*., 2012). To uncover the underlying mechanisms, we reexamined transcriptomic data from *sdg26-1* grown under normal condition (Boyu Liu *et al*., 2016) and identified several misregulated genes involved in ABA homeostasis (Supplementary Table S1). qRT-PCR analysis confirmed their misregulation (Figure 6B). Key ABA biosynthesis genes (*ABA1*, *ABA4*, *NCED5*, *NCED6*, *ABA2* and *AAO3*) were downregulated in *sdg26-1*. In line with this, the ABA exporter *ABCG25* was also reduced, whereas the ABA importer *ABCG40* and ABA-specific β-glucosidase gene *AtBG1* were upregulated. Consistent with the importance of a precise intracellular ABA homeostasis (Baron *et al*., 2012), the ABA catabolic genes *CYP707A3* and *CYP707A4* (i.e., which belong to the CYP707A subfamily, a key regulator of ABA degradation) were upregulated, particularly under non-stress conditions. Importantly, these misregulations were restored in the p+gSDG26-Myc line (Supplementary Figure 5), confirming direct and/or indirect SDG26-dependent control. Chromatin analysis provided direct evidence for this regulation at the *ABA1* locus. Indeed, H3K36me3 was lower in *sdg26-1* compared to Col0, both at control and cold conditions, and SDG26 binding was detected at the 5’ region of the gene (Figure 6C). Together, these data demonstrate that *SDG26* influence ABA homeostasis by transcriptionally regulating several genes involved in ABA biosynthesis, transport, and degradation.

ABA is a key hormone in stress adaptation (Vishwakarma *et al*., 2017). For example, it promotes stomatal closure under stress conditions, including low temperatures, thereby limiting leaf dehydration (Agurla *et al*., 2018). Given that *sdg26-1* accumulate less ABA than wild-type plants, we next assessed typical ABA-dependent physiological responses. At 21°C, the proportions of open, partially open and closed stomata were comparable among Col0, *sdg26-1*, *soc1-2* and *sdg26soc1*. Under cold, *sdg26-1* and *sdg26soc1* exhibited significantly fewer closed stomata than Col0, indicating impaired cold-induced stomatal closure (Figure 7A). Besides this, no differences in other stomatal parameters (i.e., density, index size) were detected between the different analyzed genotypes (Supplementary Figure 7). ABA is also a central regulator of plant adaptation to drought. Consistent with reduced ABA, *sdg26-1* and *sdg26soc1* plants were also more sensitive to drought stress, displaying lower survival after water deprivation and faster water loss from detached leaves (Figure 7B and C). Finally, ABA also regulates seed dormancy and germination. While all genotypes germinated similarly after stratification, *sdg26-1* and *sdg26soc1* seeds germinated more rapidly than Col0 and *soc1-2* without stratification (Figure 7D). This reduced dormancy suggests that ABA deficiency in *sdg26-1* may also extend to the embryonic stage. Together, these results show that *SDG26* is important for maintaining ABA homeostasis, which in turn influences key stress-responsive traits such as stomatal closure, drought tolerance, and seed dormancy.

**Figure 7:**
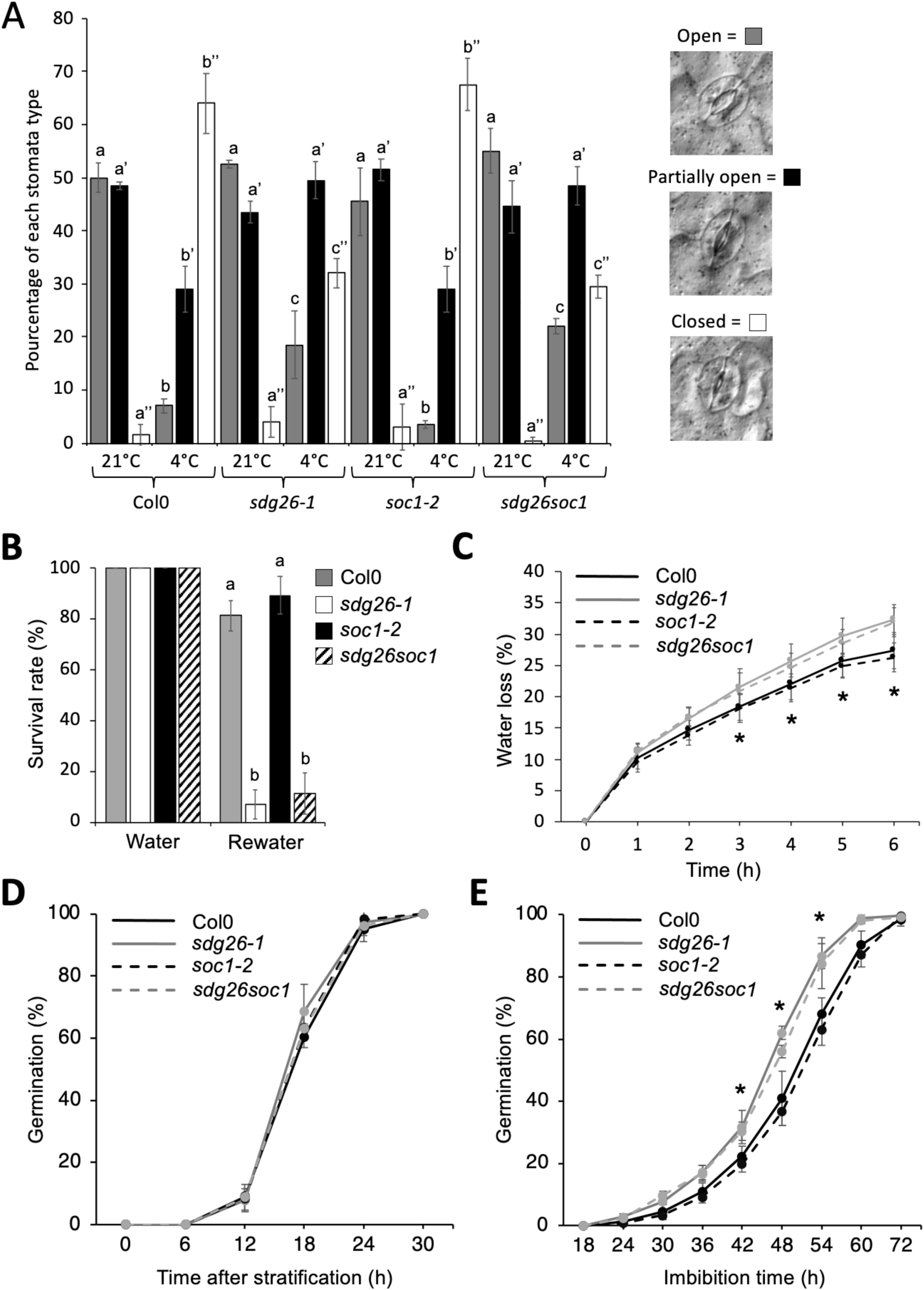
ABA-related phenotypes under cold, drought, and germination conditions. (**A**) Stomatal closure was assessed by counting the number of stomata falling into each of the three categories (Open, Partially open and Closed) before (21°C) and during a cold stress (4°C). The experiment was repeated twice with similar results. (**B**) Survival rates were after withholding water for 10 days followed by 3 days of re-watering. Data represent mean ± SD of 20 plants per genotype. The experiment was repeated three times with similar outcomes. (**C**) Water lost was quantified hourly over 6h from five detached leaves collected from three individual plants per genotypes. Data represent mean ± SD from a representative experiment. Two additional biological replicates produced similar results. Germination rates of Col0, *sdg26-1*, *soc1-2* and *sdg26soc1* seeds with (**D**) or without (**E**) stratification. Germination was scored based on radicle emergence. Data represent mean ± SD from five independent experiments with 100 seeds per genotype per replicate.

## Discussion

In this study, we provide a comprehensive characterization of the histone methyltransferase *SDG26*, demonstrating its key regulatory roles in Arabidopsis responses to abiotic stress, with a particular focus on cold stress. SDG26 was originally classified as a TrxG histone methyltransferase based on its SET domain and phylogenetic similarity to ASH1-type enzymes (Baumbusch *et al*., 2001). Consistent with other TrxG proteins (Zhao *et al*., 2005)(Puig *et al*., 2007)(Berr *et al*., 2009)(Chen *et al*., 2017), SDG26 localizes to the nucleus and shows widespread expression across organs, (Figure 2). *In situ* hybridization revealed transcript enrichment in young tissues and reproductive structures such as anthers and ovules (Figure 6), echoing the expression domains of *SDG8* and *SDG2* (Zhao *et al*., 2005)(Berr, McCallum, Ménard, *et al*., 2010). Promoter-GUS analyses further reveal both overlapping and distinct expression domains among TrxG proteins. Similar to *ATX1-5* (Saleh *et al*., 2007)(Hayot *et al*., 2012)(Chen *et al*., 2017), the *SDG26* promoter drives strong GUS activity in vascular tissues (Figure 7), a pattern absents from *SDG7* and *SDG8* promoters (Kim *et al*., 2005)(Berr, McCallum, Alioua, *et al*., 2010) (Lee *et al*., 2015). In root tips, GUS staining was detected for *ATX1, ATX2, SDG7, SDG8*, and *SDG26* (Saleh *et al*., 2007)(Berr, McCallum, Alioua, *et al*., 2010)(Lee *et al*., 2015)(Figure 7E), but not for *ATX3-5* (Chen *et al*., 2017). Likewise, GUS activity in flowers is broadly shared among *SDG26*, *SDG8*, and *ATX1-5* (Saleh *et al*., 2007)(Cazzonelli *et al*., 2009)(Hayot *et al*., 2012)(Chen *et al*., 2017)(Figure 7D). Notable differences have also been reported, including *SDG8* activity in hydathodes (Berr, McCallum, Alioua, *et al*., 2010) and *ATX3-5* expression in trichomes (Chen *et al*., 2017). Collectively, these observations show that TrxG genes display broad but also partially overlapping tissue-specific expression consistent with the physical interactions already documented between SDG26 and ATX1 or SDG8 (Valencia-Morales *et al*., 2012). Together, these results suggest that, in connection or not with its function in flowering-time regulation, SDG26 is likely involved in additional developmental processes.

In this work, we demonstrated that SDG26 is induced by abiotic stresses but not by stress-related phytohormones (Figure 9), a regulation confirmed at both transcript and protein levels (Figure 10) and apparently conserved, since the SDG26 ortholog in *Populus trichocarpa* is also strongly upregulated in dormant winter stems (Ko *et al*., 2011). Although transcript and protein levels were stress-responsive, SDG26 nuclear localization remained stable, with no evidence for long-distance mobility or altered subnuclear distribution under stress (Figures 8, 10). Previously, we established that SDG26 regulates *SOC1* during flowering (Berr *et al*., 2015)(Boyu Liu *et al*., 2016) and *SOC1* itself was proposed to mediate crosstalk between cold sensing and flowering (Seo *et al*., 2009). Here, we observed that *SOC1* was rapidly induced by cold in Col0 plants (Figure 4A), whereas its induction was reduced in *sdg26* and correlated with lower enrichment of H3K4me3 and H3K36me3 at *SOC1* chromatin (Figure 4A-F). Despite this induction, SDG26 occupancy at *SOC1* chromatin in Col0 remained stable during cold treatment, suggesting that its contribution to *SOC1* activation does not involve dynamic recruitment but rather the maintenance of permissive chromatin states (Figure 4E-F). Seo et al. (2009) proposed that *SOC1* represses *CBF-COR* genes. Yet, despite reduced *SOC1* induction in *sdg26*, *CBF-COR* genes were also less induced by cold (Figure 5A). This result is consistent with our finding that SDG26 directly binds *CBF* loci, where its occupancy and associated H3K36me3 enrichment increase after cold exposure but are reduced in *sdg26* (Figure 5B–E). Together these findings indicate that SDG26 contributes to cold responses not only through *SOC1* but also via activation of *CBF* genes. This dual role raises the possibility that SOC1 and SDG26 may compete or cooperate for binding at *CBF*, possibly through shared cis-regulatory elements. In addition, since CBF expression is modulated by transcription factors such as ICE1/2, CAMTA1–3, PIF4/7, and MYB15 (Agarwal *et al*., 2006)(Doherty *et al*., 2009)(Lee and Thomashow, 2012), SDG26 recruitment may also be coordinated with these factors. Clarifying whether SDG26 acts alone or in combination with such regulators remains an open question.

Beyond its role in *SOC1* and *CBF* regulation, our results show that SDG26 also contributes to cold responses through ABA-dependent pathways by maintaining ABA homeostasis, thereby supporting ABA-mediated processes such as stomatal regulation, drought tolerance, and seed dormancy. Among the downregulated genes in *sdg26*, *ABA1* and *ABA2* are known to promote flowering (Riboni *et al*., 2016), suggesting that the late-flowering phenotype of *sdg26* may partly result from reduced endogenous ABA. Interestingly, the rice homolog of *SDG26*, *SDG708*, also promotes flowering, and its loss of function results in delayed flowering similar to Arabidopsis *sdg26* mutants (Sui *et al*., 2013), pointing to a conserved role of SDG26-like proteins in reproductive timing. Since ABA promotes early flowering in rice (Du *et al*., 2018), it will be important to determine whether SDG708, like Arabidopsis SDG26, also contributes to ABA homeostasis in a conserved mechanism linking chromatin regulation, ABA signaling, and flowering across species. Additional pathways may further connect ABA signaling to *SOC1* regulation. Under drought stress, ABF3 and ABF4 together with NF-YCs were shown to promote *SOC1* transcription, thereby accelerating flowering and facilitating life cycle completion under stress (Hwang *et al*., 2019). This raises the possibility that SDG26 influences flowering and stress adaptation both through direct chromatin regulation of *SOC1* and indirectly by modulating ABA levels, which could feed into ABA-dependent *SOC1* induction under specific stress conditions. Finally, ABA itself has long been recognized as a central regulator of cold acclimation (Shi *et al*., 2015), reinforcing the view that SDG26 integrates ABA-dependent and ABA-independent pathways to fine-tune the balance between stress resilience and reproductive development. SDG26 thus sits at a critical crossroads between flowering regulation, ABA signaling, and abiotic stress response. Its known role in activating SOC1 and modulating flowering time, combined with its effect on ABA levels and stress tolerance, supports a model in which SDG26 orchestrates a coherent developmental and physiological program. The combination of late flowering, reduced ABA accumulation, impaired stomatal dynamics, enhanced freezing tolerance, and drought susceptibility observed in *sdg26* plants is consistent with ABA’s dual role in promoting both stress resilience and reproductive transition (Jiang *et al*., 2025). As such, SDG26 may help balance these processes under adverse conditions, allowing plants to optimize survival while ensuring timely flowering, and potentially promoting flowering as a strategy to complete the life cycle under stress. This dual positioning is not unprecedented. ATX1 regulates drought responses via an ABA-dependent pathway (Ding *et al*., 2011), and protein–protein interactions between ATX1 and SDG26 have been reported (Valencia-Morales *et al*., 2012), raising the possibility of cooperative regulation of ABA-related stress responses. Similarly, SDG8 and ATX1 are also known to regulate both flowering and stress pathways (Ramirez-Prado *et al*., 2018). Such integrated roles may provide an adaptive advantage by coordinating development and stress acclimation to maximize survival and reproductive success under fluctuating environments.

Finally, functional studies have shown that SDG26 promotes H3K4 and H3K36 trimethylation at target loci such as *SOC1*, thereby impacting transcription (Berr *et al*., 2015)(Boyu Liu *et al*., 2016). More recent evidence indicates that SDG26 can also participate in repressive complexes, as in the case of *FLC* regulation where it associates with RNA-processing factors and influences H3K27me3 states (Fang *et al*., 2020)(Qi *et al*., 2022). Our results extend this view by showing that SDG26 directly binds cold-responsive loci, including *CBF* genes and *ABA1*, and that its loss reduces H3K36me3 enrichment at these sites, underscoring its versatile role in gene regulation. This versatility reflects a broader paradigm in plant chromatin, where histone methylation, and in particular H3K36, acts as a central regulator of stress responses by shaping both acute transcriptional reprogramming and transcriptional memory (Yu *et al*., 2025). A recent study showed that SDG26 plays a context-dependent and cooperative role in fine-tuning chromatin states by contributing to H3K36me3 deposition. Together with SDG8 and SDG25, it helps establish locus-specific H3K36me3 landscapes, including at long, stress-responsive genes, thereby contributing to the formation of a “transcription-resistive” chromatin state, an open but transcriptionally inert configuration associated with transcript stabilization (Yao *et al*., 2025). Within this framework, SDG26 emerges as a context-dependent and cooperative factor that fine-tuning chromatin states at selected genes, thereby integrating developmental and environmental cues into adaptive transcriptional programs.

## Author contribution

A. B. initiated and coordinated the project. A. B. and X. Z. designed the experiments. X. Z., J. Z., M. E., and A. B. performed experiments. A. B. and X. Z. made figures and wrote the manuscript. J. Z., M. E. and W-H. S. revised the manuscript. All authors approved the final version of the manuscript.

## Declaration of interest

The authors declare no competing interests.

## Notes

### Competing Interest Statement

The authors have declared no competing interest.

