## Supplementary figures and images for "The histone methyltransferase SDG26 shapes cold stress responses in Arabidopsis through chromatin-based regulation of ABA-dependent and ABA-independent pathways"

### Supp Figures

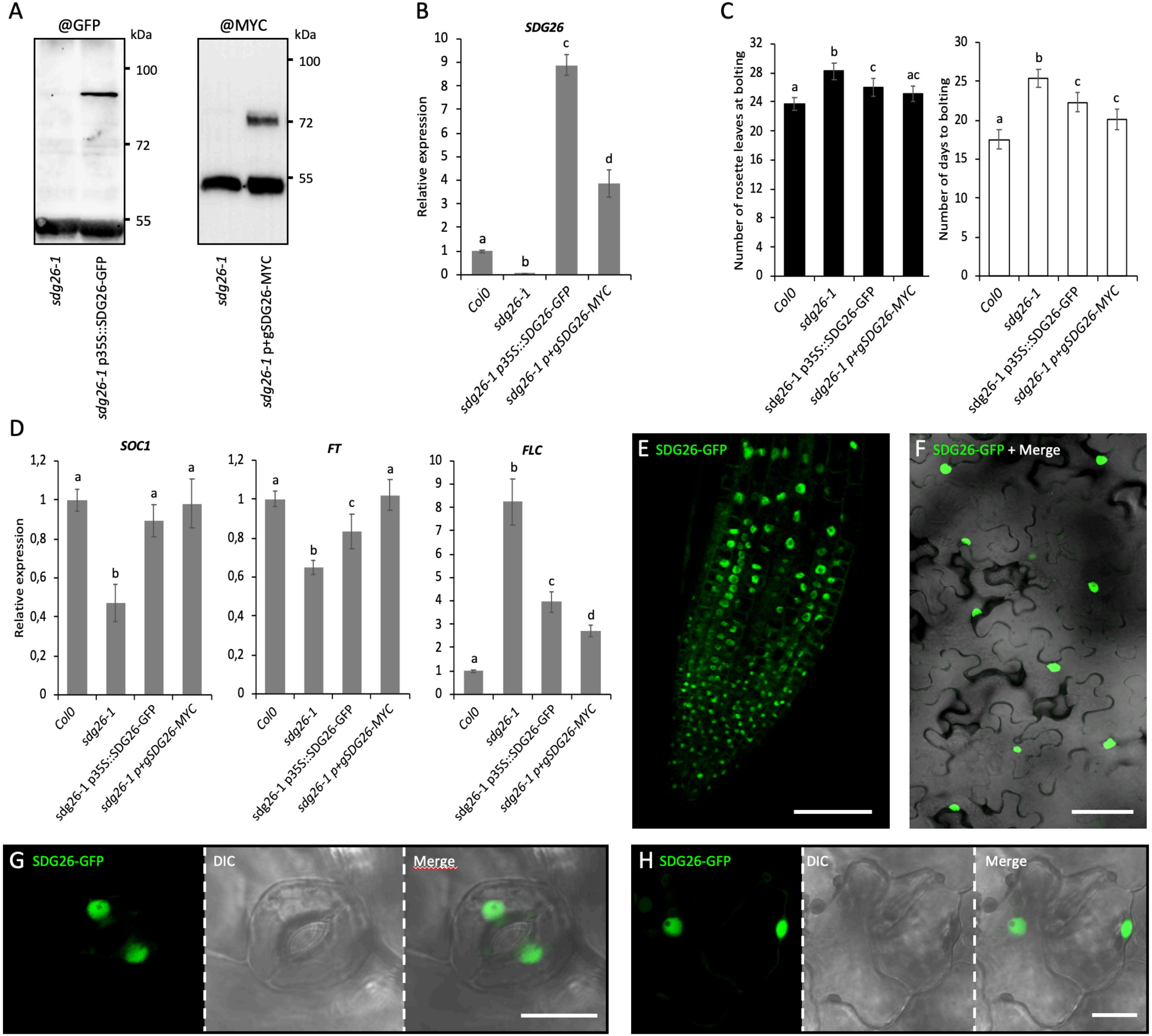

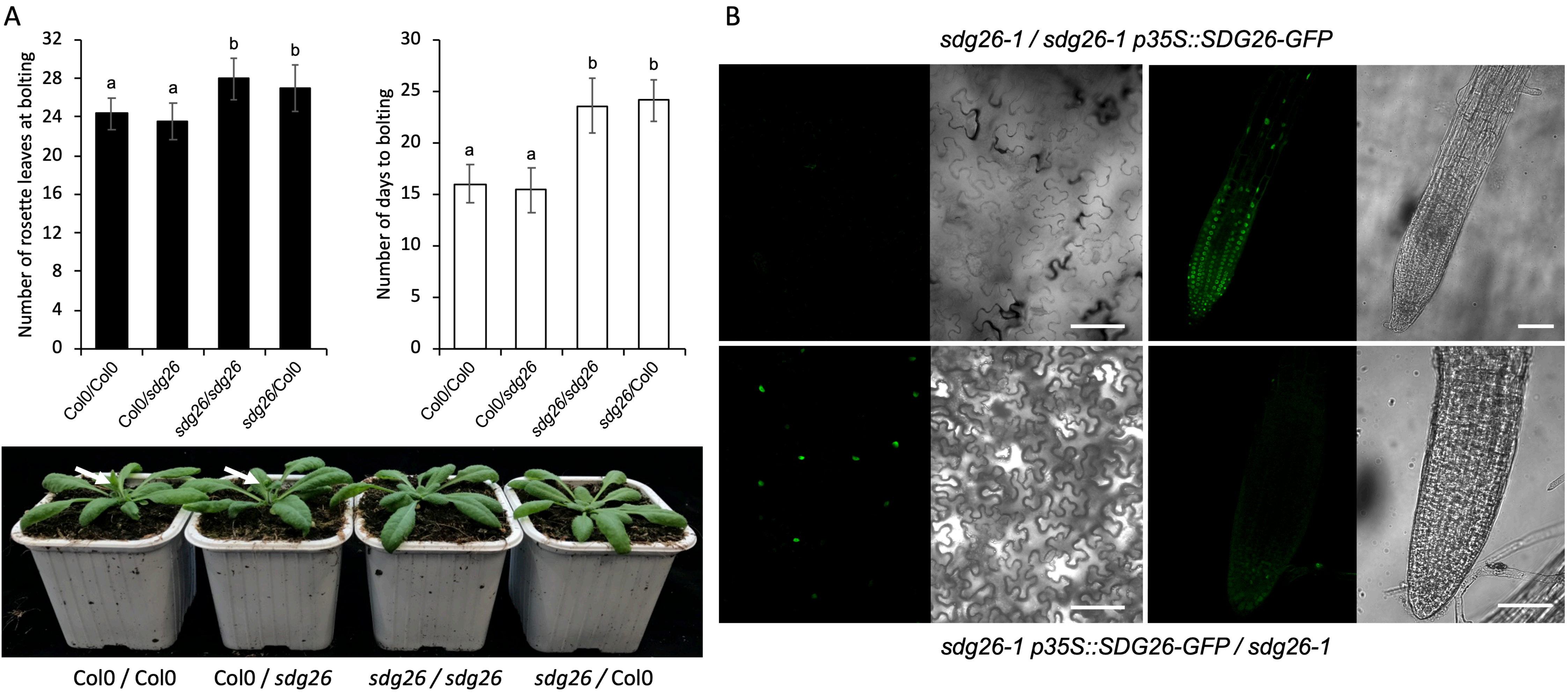

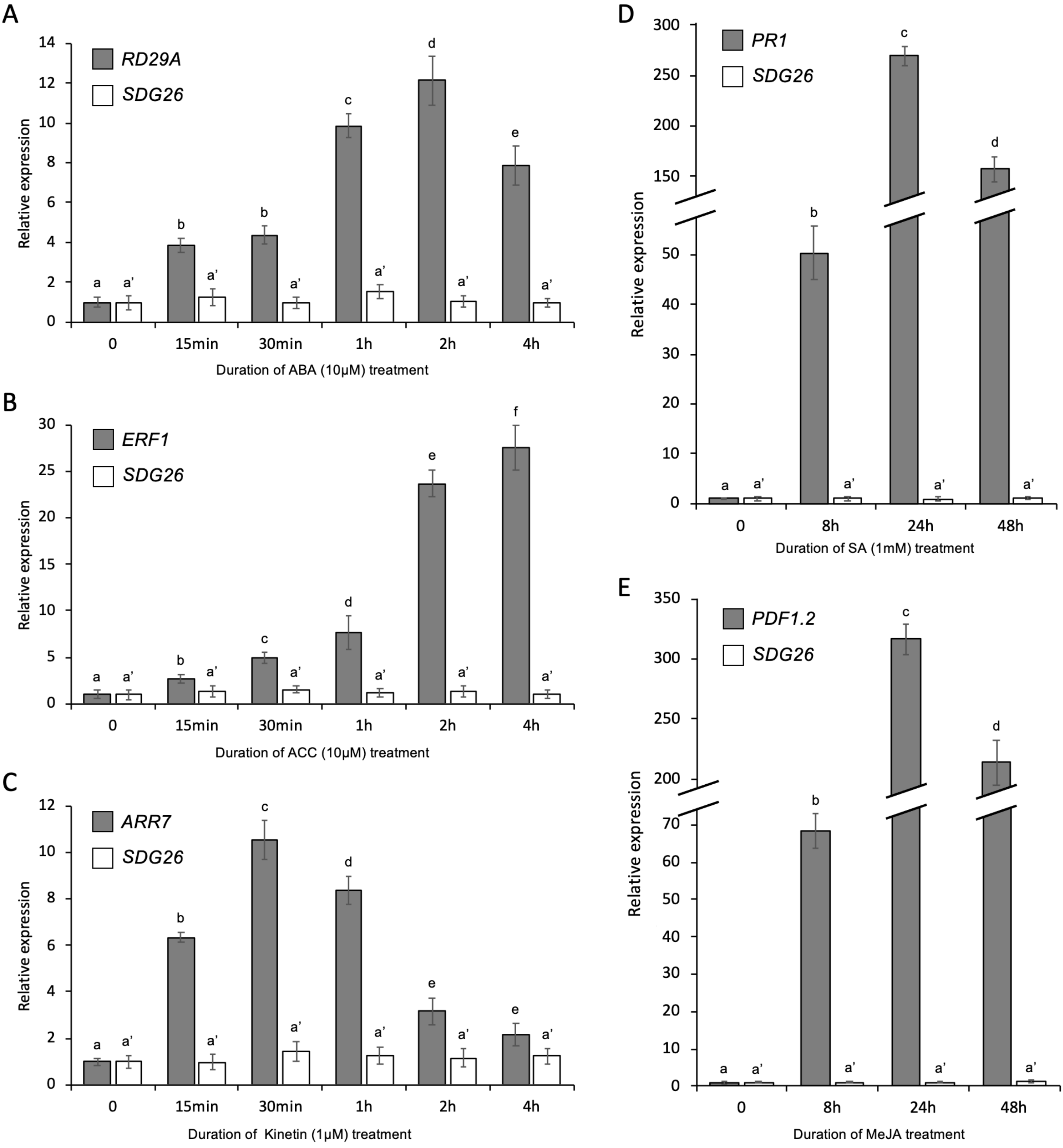

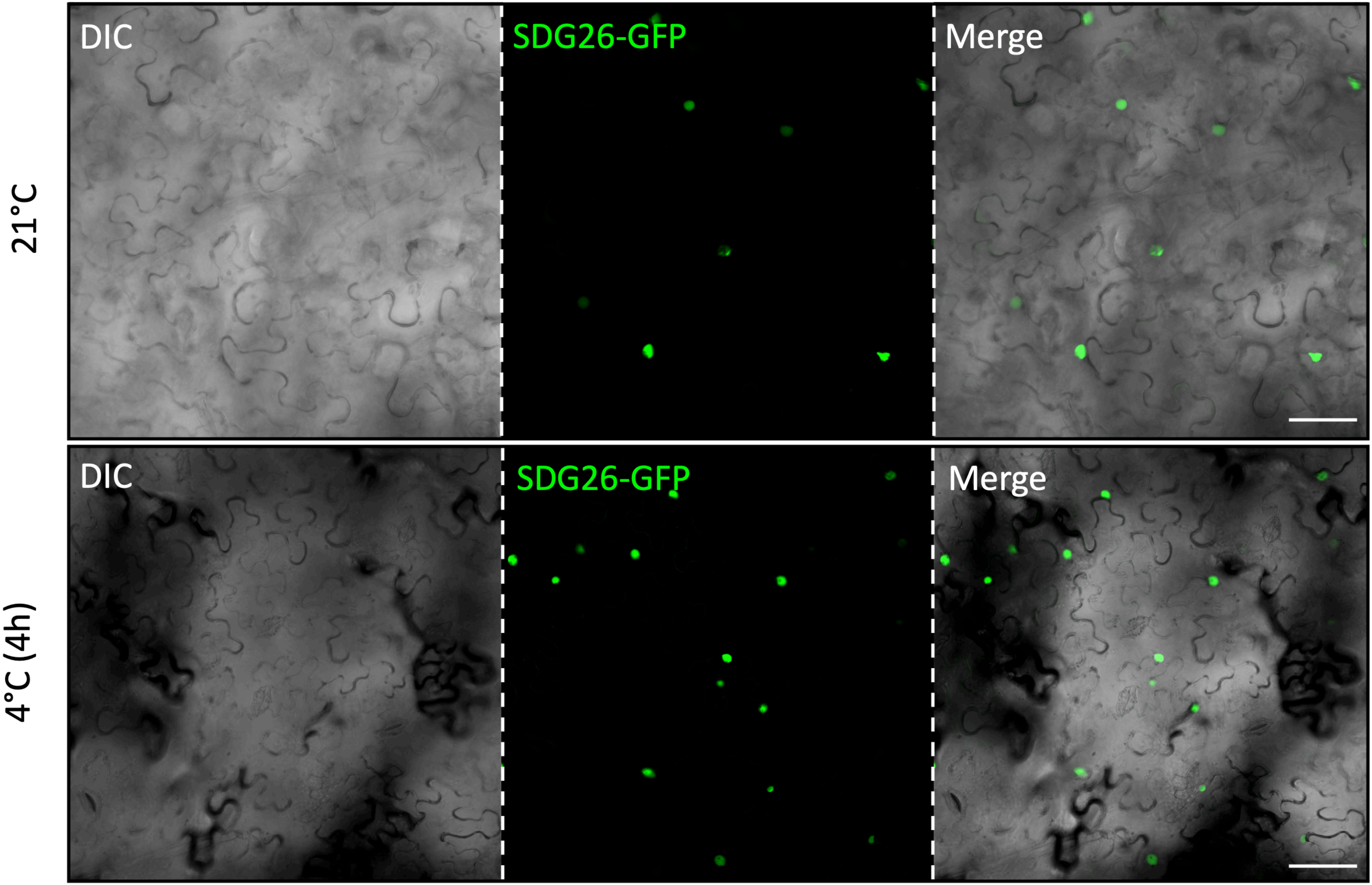

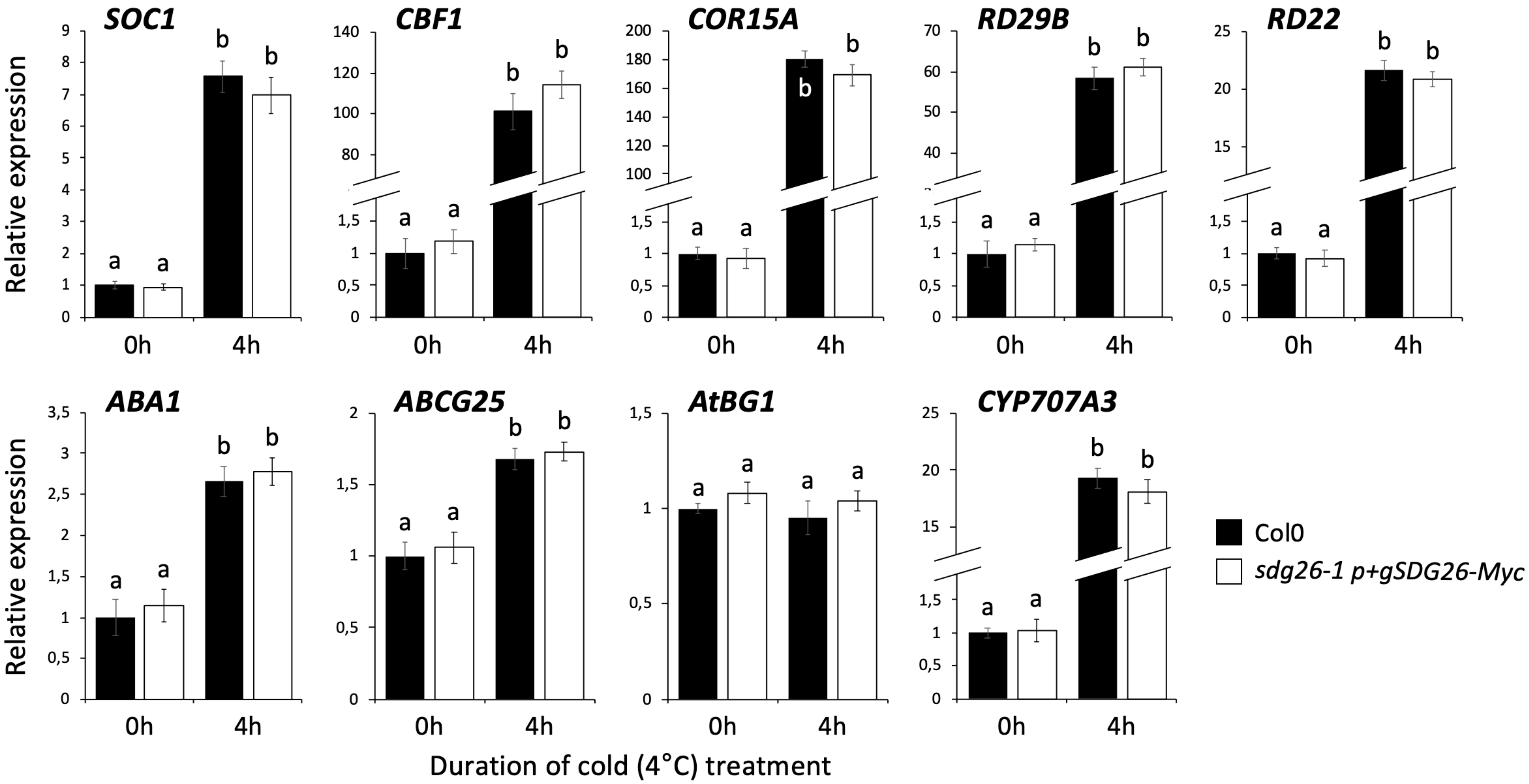

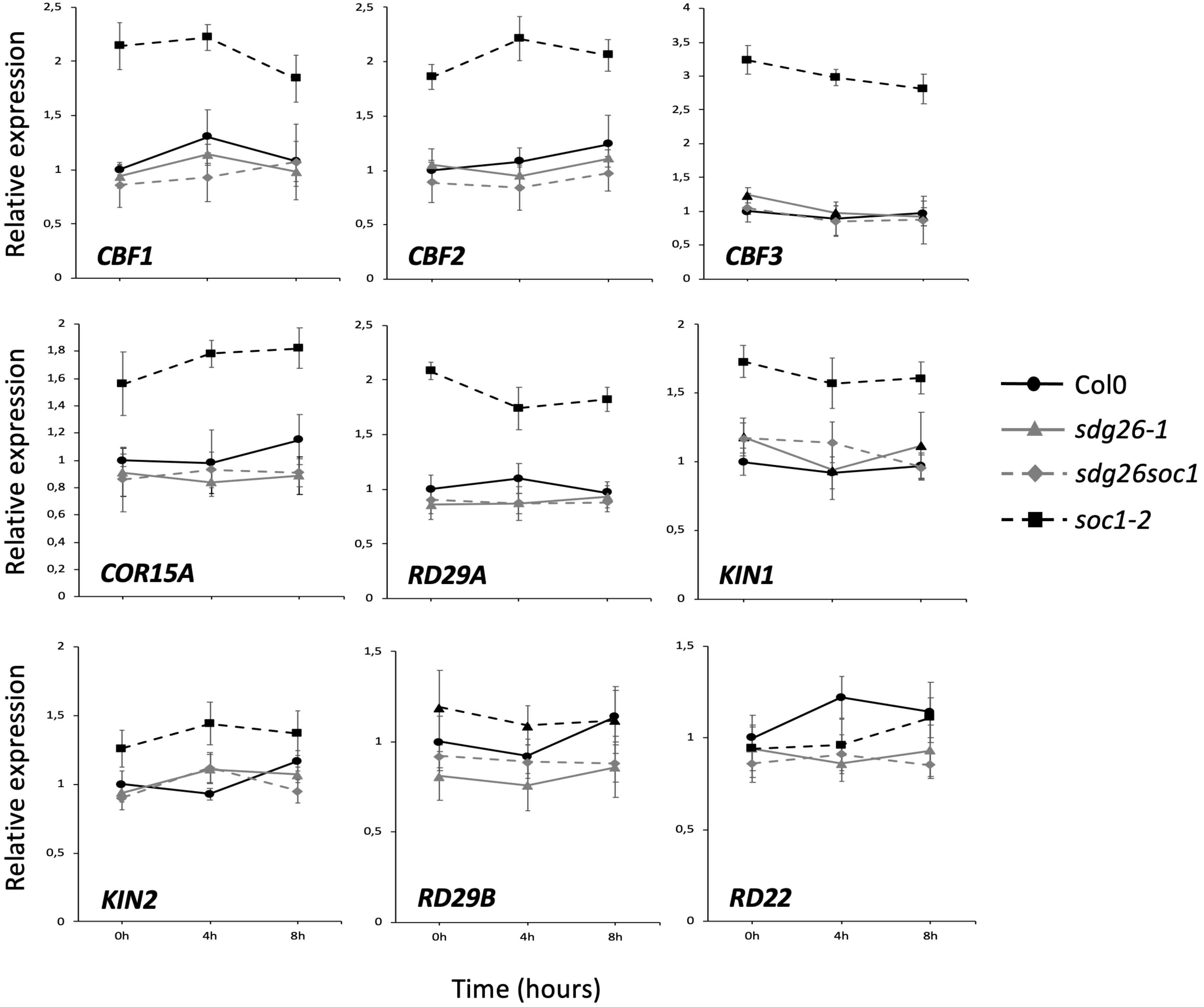

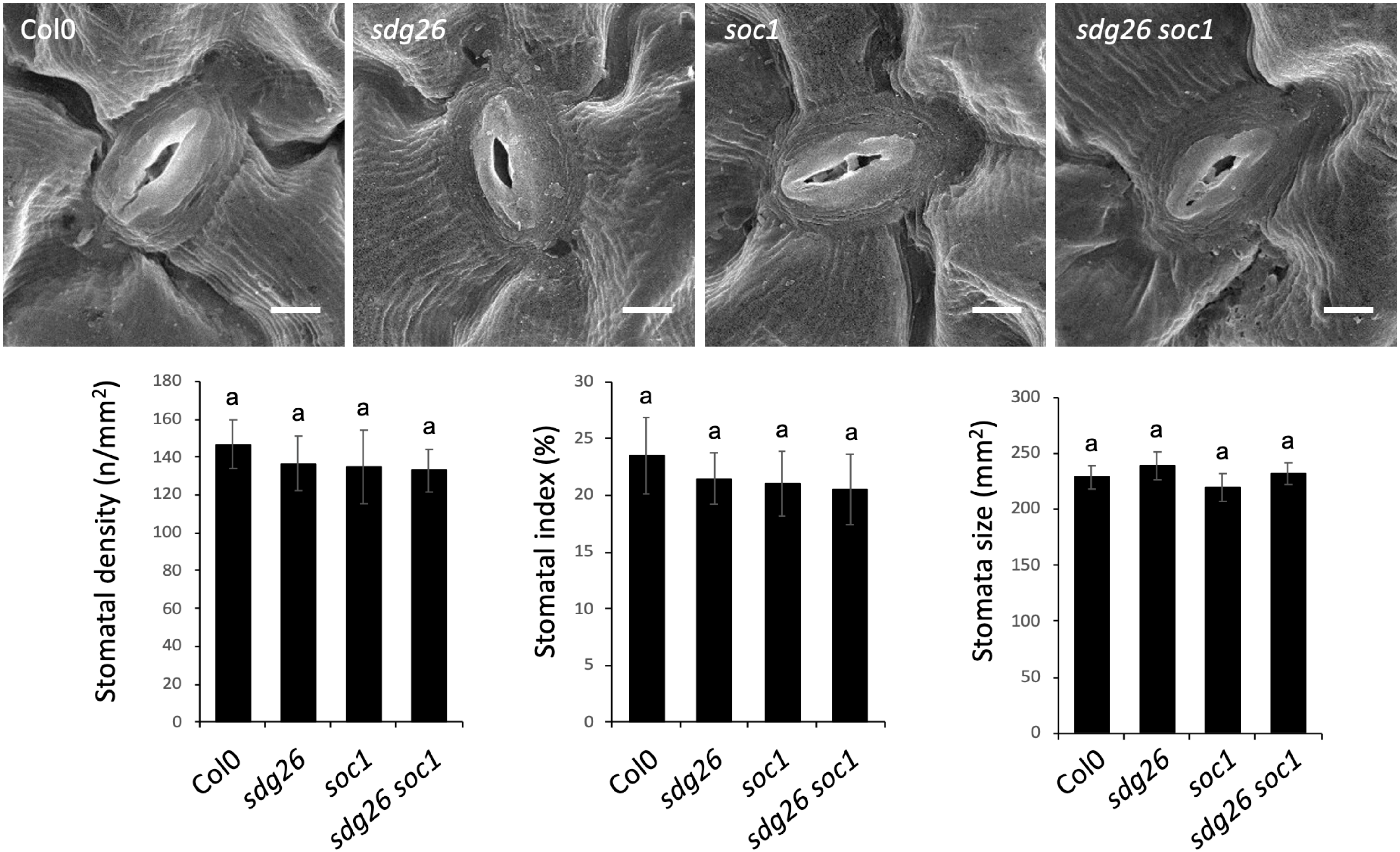

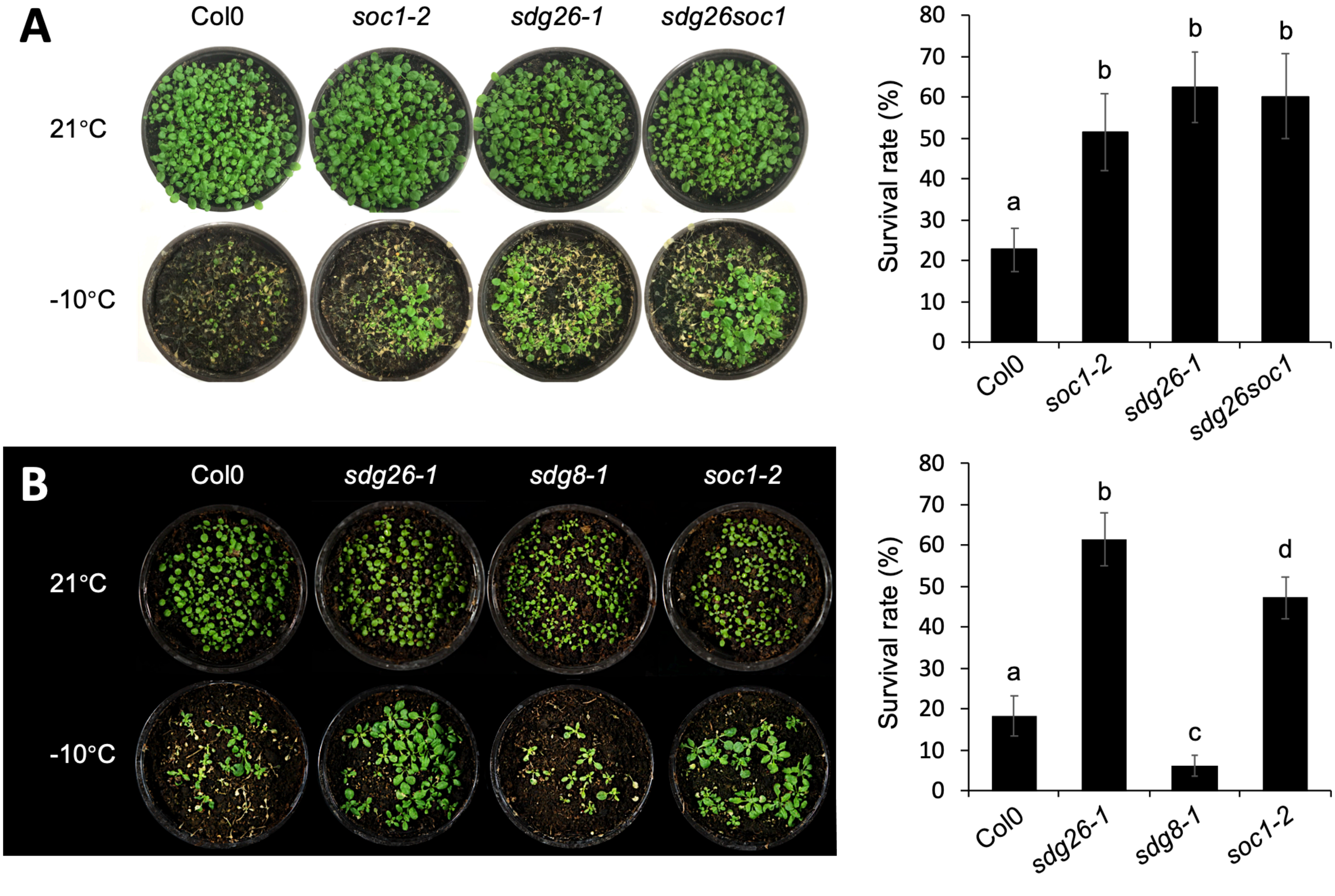
